# An updated suite of viral vectors for *in vivo* calcium imaging using local and retro-orbital injections

**DOI:** 10.1101/2021.05.14.443815

**Authors:** Sverre Grødem, Ingeborg Nymoen, Guro Helén Vatne, Valgerdur Björnsdottir, Kristian Kinden Lensjø, Marianne Fyhn

## Abstract

Calcium imaging using genetically encoded Ca^2+^ indicators (GECIs) is a widely adopted method to measure neural activity in modern neuroscience. Here, we explore the use of systemically administered viral vectors for brain-wide expression of GECIs, and adapt novel GECIs to optimize signal-to-noise. We show that systemic injections of PHP.eB AAVs to express GECIs is a highly promising technique for imaging neural activity and circumvent the need for transgenic GECI expressing mouse lines. We also establish the use of soma-targeted GECIs that outperform current Ca^2+^ indicators using both systemic and local virus injections.

## INTRODUCTION

The use of microscopy to measure the activity of neurons is widely applied in modern neuroscience. With the development of genetically encoded Ca^2+^ indicators (GECIs) there has been rapid advances in response kinetics, sensitivity and brightness of Ca^2+^ sensors (e.g.^1,2^), of which the GCaMP sensors are the most prominent. These engineered proteins contain a Ca^2+^ binding motif and a circularly permuted green fluorescent protein that brightens when Ca^2+^ is present. Using GECIs for activity measurements allows for cell-type targeted recordings, repeated measurements of the same cells for up to several months and recordings from large populations of neurons. Ideally, GECIs should be uniformly expressed across the neuronal population. Overexpression in a subset of cells can lead to intracellular aggregation and eventually cell death.

To achieve this, various methods to introduce the genetic constructs into cells have been explored. An often preferred method for introducing GCaMP into neurons is by way of transgenic animal models(e.g.^3,4,5^). These models allow for strong, even and sustained expression throughout life that can be targeted to specific cell types. However, transgenic GCaMP mice require intricate breeding schemes that come with high costs, both financial and for animal welfare. They also depend on driver lines that prevent the use of other transgenics^3^. Moreover, because the GECIs are expressed throughout development, frequent ictal activity has been reported for several such mouse strains^6^, questioning their reliability.

GCaMP can also be delivered to cells by a viral vector, either through a local injection directly into the tissue of interest (e.g.^7^) or by intracerebroventricular injections^8^. Furthermore, using adeno-associated virus (AAV) serotype 9 which crosses the blood-brain-barrier^9^ in neonatal mice, GCaMP may be introduced through an intravenous injection into the tail vein, temporal vein^10^ or transverse sinus^11^. However, these administration techniques are technically challenging. They also come with the risk of overexpression of the GECI because of the young age at the time of injection, and extended period from injections to experiments which may lead to cell damage or ictal events. Local injections of viral vectors tend to lead to highly variable expression depending on the concentration of virus particles and is often associated with cell damage or death^12^.

In contrast to AAV9, the recently developed AAV serotype PHP.eB has been shown to cross the blood-brain barrier in adult animals and efficiently transduce neurons across the brain^13^, suggesting that genes could be delivered via intravenous injections^14^. Importantly, such injections can be performed at any stage in development and thus prevent accumulation of GCaMP and disturbing Ca^2+^ homeostasis during sensitive parts of development. Moreover, this would enable brain-wide expression of the GECI in combination with other transgenic models for e.g. cell-type specific activity perturbations. In contrast to tail vein and other intravenous injection procedures, injections into the retro orbital (RO) sinus can be performed with minimal training. RO injections are quick, non-invasive, and impose little stress to the animals compared to other methods^15^.

Here we present a GECI screening in mice applying the RO injection method for systemic viral delivery and assess functionality using widefield and two-photon laser-scanning microscopy. All viral vectors were tested by both RO and local injections to verify the efficiency of the constructs. Reduced neuropil contamination is favorable for population imaging. Thus, we restricted expression to the cell soma utilizing two soma-targeting approaches^16,17^. Both EE-RR soma-targeted and ribosome-tethered (RL10) jGCaMP8 resulted in reduced neuropil contamination and the latter led to highly selective expression in the cell soma. For local injections, ribosome-tethered GECIs showed very high density of cell labeling, but came with a long delay in expression. To circumvent this issue we introduce a modified ribosome-tethered GECI that shows strong expression just one week after local injections. In general, we observe a remarkable signal-to-noise ratio using both systemic and local virus administration, enabling activity detection from a higher number of cells as their activity is not masked by the surrounding neuropil.

We show that several recently developed GECIs are highly suitable for this application and give rise to uniform and stable expression for many weeks and can be combined with other transgenic models for e.g. cell-specific expression of optogenetic or chemogenetic receptors. Finally, we combine modern constructs for restricting GCaMP localization to the cell soma^16,17^ with the most recent iterations of jGCaMP, the jGCaMP8^18^ sensors. With these soma-targeted GECIs we observe remarkable signal-to-noise ratio, using both systemic or local virus administration.

## RESULTS

### Performance screening of existing GECIs using RO injections

We initially screened the performance of 10 existing GECI variants administered by RO or local injections in high titre PHP.eB serotype AAVs. In addition, two fluorescent proteins were expressed by RO and local PHP.eB injections (mNeonGreen expressed under a CAG promoter, and floxed tdTomato was injected in PV-Cre mice). Briefly, pairs of mice were randomly assigned a GECI-expressing AAV and evaluated every two weeks for 2,5 months using widefield and two-photon imaging through a cranial window, followed by post-mortem histology (**Fig. 1a, S1, S2**). The majority of screened GECIs, which are variations of GCaMP, were not sufficiently bright for in-vivo calcium imaging following systemic virus administration (**Fig. 1, Table 1**). The most widely used GECI, GCaMP6f was undetectable at reasonable laser power when expressed from an intravenously injected AAV, likely due to reduced multiplicity of infection (MOI) associated with intravenously administered viruses. Notably, GCaMP6f was present and visible in the tissue after 6 weeks, but only when using very high laser power (>140 mW output at the objective) which would not be sustainable for functional experiments (**Fig. 2a**). In an attempt to improve the brightness, we tested both double and triple injection volumes of RO administered GCaMP6f, but the resulting expression was still too dim to image at reasonable laser power (data not shown).

**Table 1:** Overview of initial GECI screening with virus titer and Addgene reference indicated.

| GECI | Titre (VG/ml) | Brightness at 50mW (RO injection) | Neuropil | Brightness at 50mW (local injection) | Addgene plasmid # |
| --- | --- | --- | --- | --- | --- |
| GCaMP6f | 2.44E+13 | none | NA | medium | 100837 |
| Soma-GCaMP6f | 2.04E+13 | low | low | medium | 158756 |
| jGCaMP7f | 2.04E+13 | low | NA | medium | 104488 |
| Soma-GCaMP7f | 1.53E+13 | low | low | medium | 158760 |
| jGCaMP7s | 1.11E+13 | high | medium | high | 104487 |
| jGCaMP8f | 1.04E+13 | low | high | medium | 162376 |
| jGCaMP8m | 9.43E+12 | medium | high | high | 162375 |
| jGCaMP8s | 1.29E+13 | high | high | high | 162374 |
| JREX-GECO1 | 9.36E+12 | high | low | high | 169259* |
| Ribo-GCaMP6m | 1.86E+13 | low | low | low | 158777 |
| CAG-mNeonGreen | 1.09E+13 | high | NA | high | 99134 |
| TdTomato # | 8.83E+12 | high | NA | NA | 28306 |

**Table 2:**
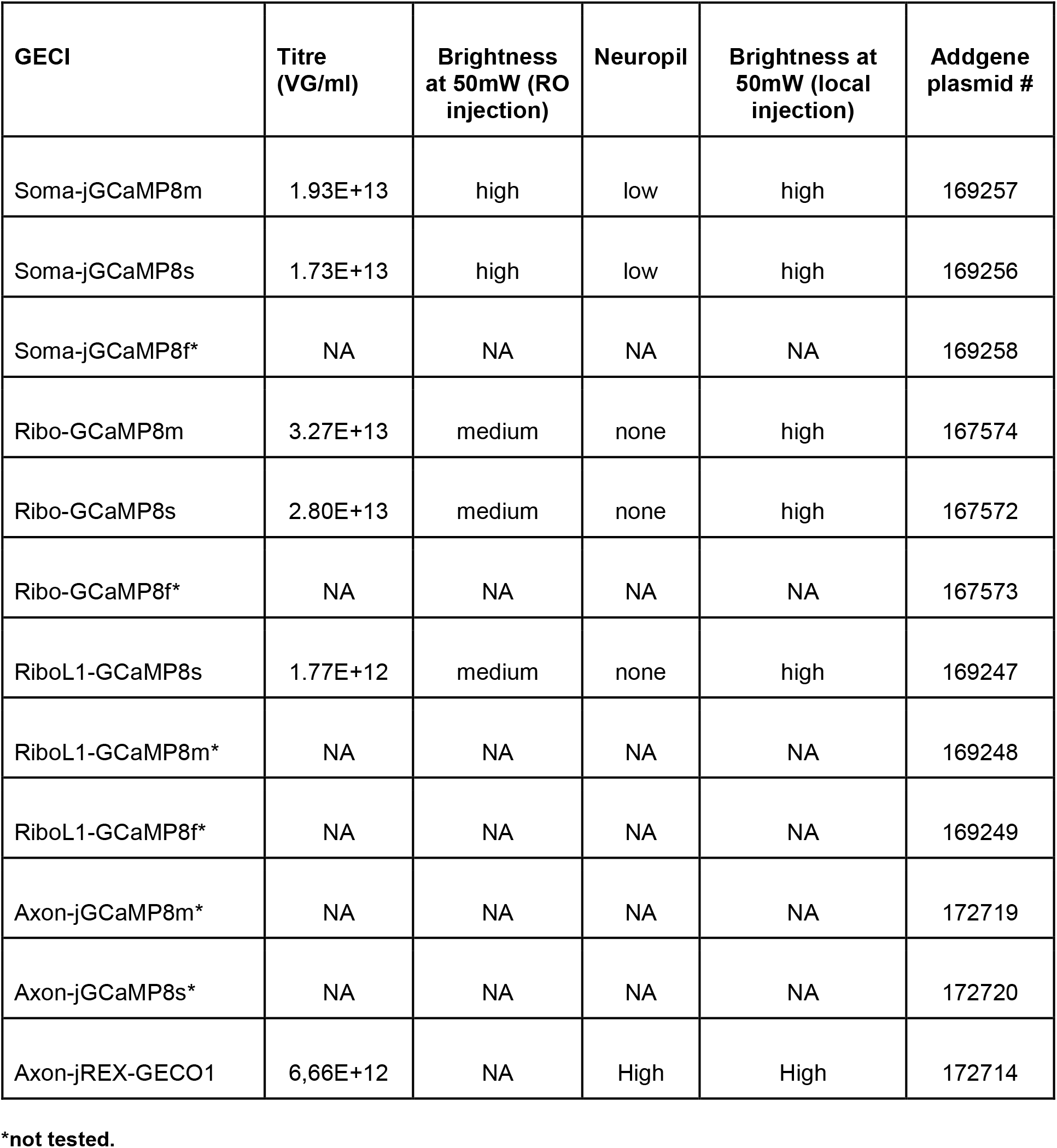
Soma and axon-targeted GECI constructs made in this work.

| GECI | Titre (VG/ml) | Brightness at 50mW (RO injection) | Neuropil | Brightness at 50mW (local injection) | Addgene plasmid # |
| --- | --- | --- | --- | --- | --- |
| Soma-jGCaMP8m | 1.93E+13 | high | low | high | 169257 |
| Soma-jGCaMP8s | 1.73E+13 | high | low | high | 169256 |
| Soma-jGCaMP8f* | NA | NA | NA | NA | 169258 |
| Ribo-GCaMP8m | 3.27E+13 | medium | none | high | 167574 |
| Ribo-GCaMP8s | 2.80E+13 | medium | none | high | 167572 |
| Ribo-GCaMP8f* | NA | NA | NA | NA | 167573 |
| RiboL1-GCaMP8s | 1.77E+12 | medium | none | high | 169247 |
| RiboL1-GCaMP8m* | NA | NA | NA | NA | 169248 |
| RiboL1-GCaMP8f* | NA | NA | NA | NA | 169249 |
| Axon-jGCaMP8m* | NA | NA | NA | NA | 172719 |
| Axon-jGCaMP8s* | NA | NA | NA | NA | 172720 |
| Axon-jREX-GECO1 | 6,66E+12 | NA | High | High | 172714 |

\*not tested.

**Figure 1:**
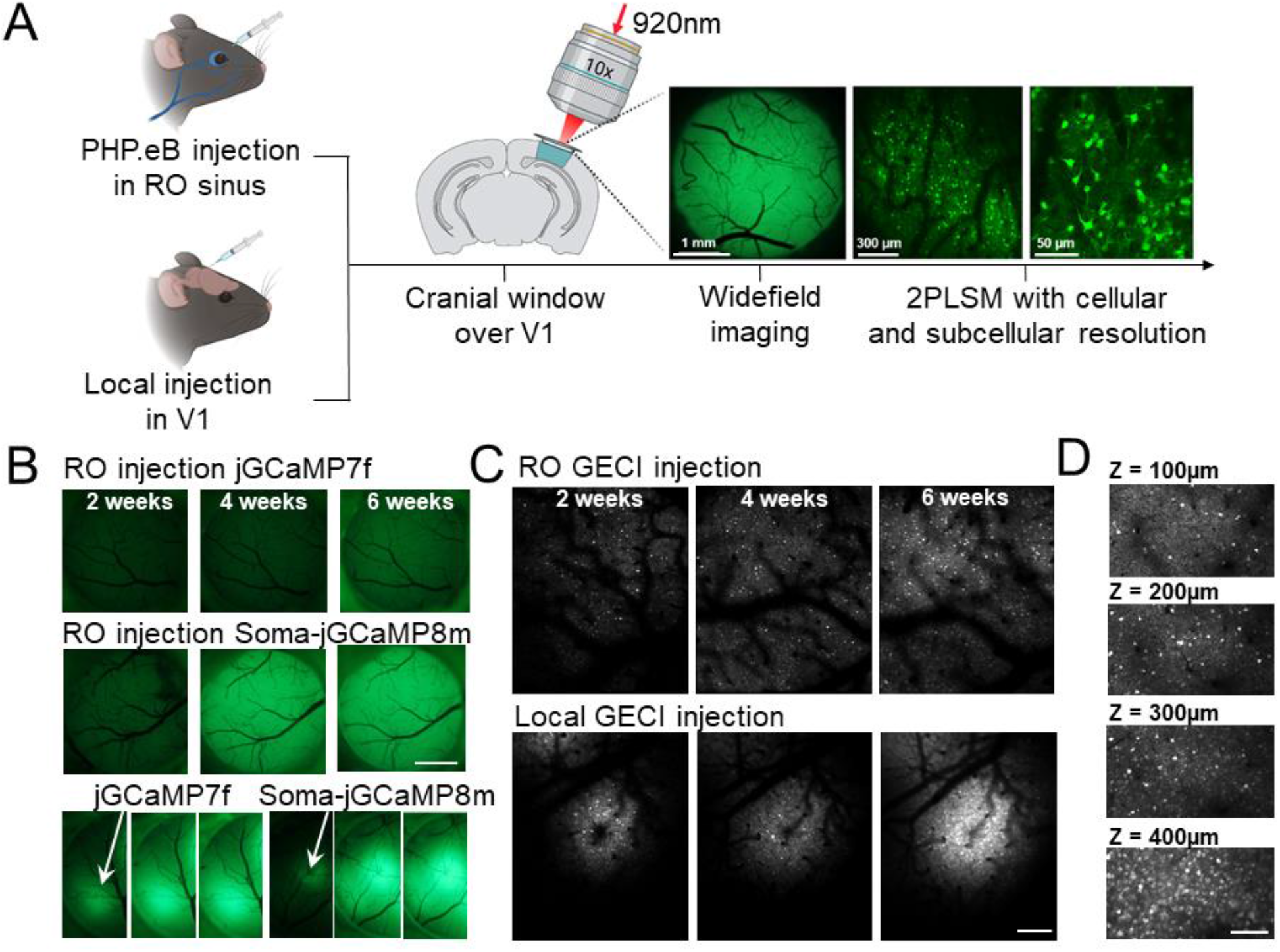
GECI screening after local or retro-orbital (RO) virus injections. **A**: Experimental overview indicating the two injection approaches, and the methods used to monitor the expression. **B**: Example images from widefield fluorescence microscopy of two GECIs expressed by RO (two upper panels) or local (lower panel) virus injections 2, 4 and 6 weeks after injection. Scale bar indicates 1 mm. **C**: Example images from *in vivo* two-photon microscopy of a GECI expressed by RO or local virus injection. Scale bar indicates 250 μm. **D**: GECI expression at different depths in cortex after RO injection. Scale bar indicates 150μm.

**Figure 2:**
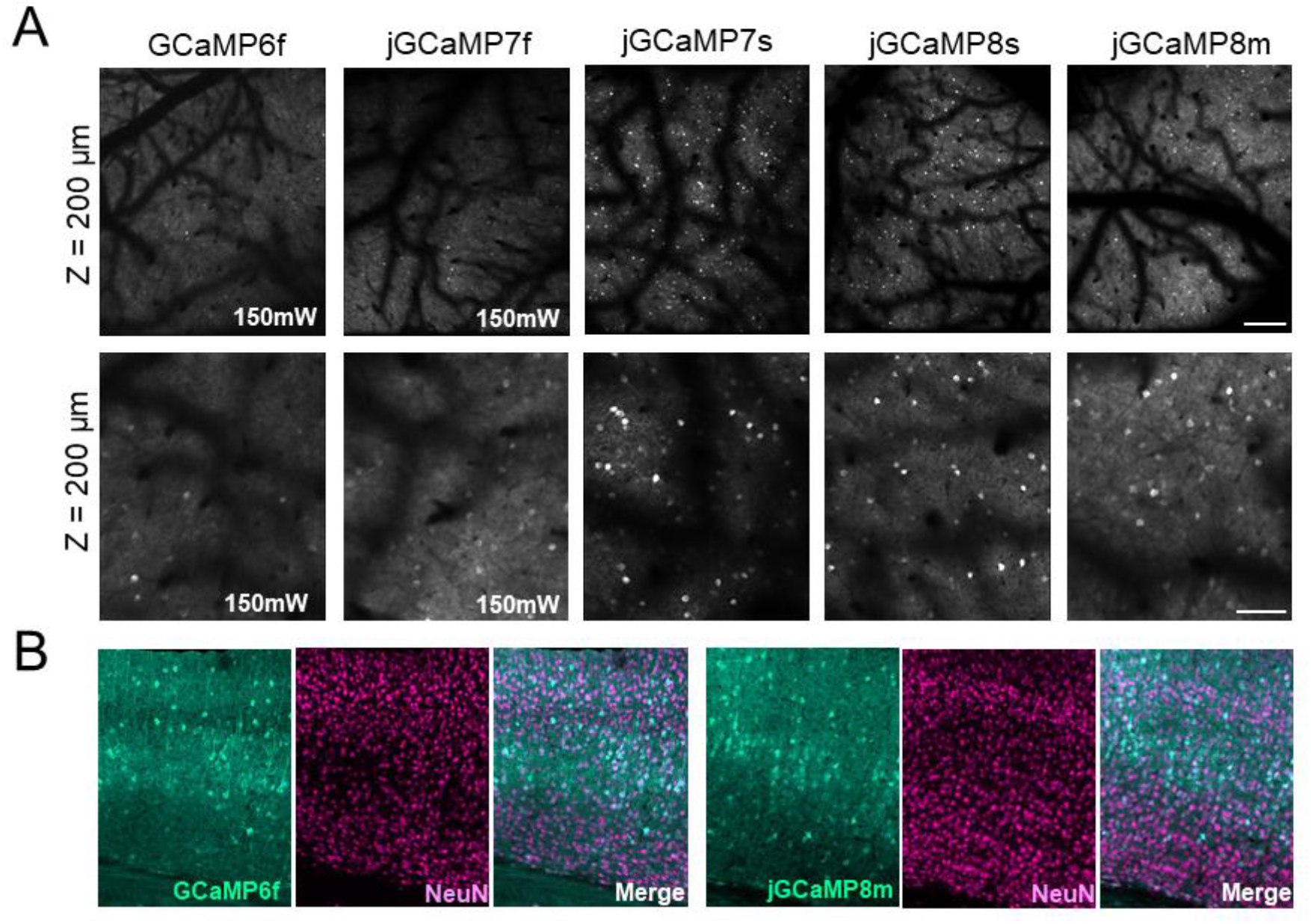
*In vivo* GECI screening after retro-orbital virus injections. **A**: Example images from five different GECIs at different magnification. Note that GCaMP6f and jGCaMP7f were not detectable at reasonable illumination intensity (40-50mW at 920 nm). The images shown from GCamp6f and jGCaMP7f were acquired using 150mW laser power for testing purposes only. The recently developed jGCaMP7s, 8s and 8m were sufficiently bright and were imaged using 50mW. Scale bar indicates 250 μm (upper panel) and 100μm (lower panel). All images shown are average intensity projections from 2000 frames with identical adjustments to brightness and contrast. **B**: Histology images from primary visual cortex six weeks after RO injection. The expression of GCaMP6f and jGCaMP8m were similar, showing strong expression throughout the cortex, in particular in layer 5 neurons.

Of the more recently developed GECIs; jGCaMP7s, jGCaMP8s and jGCaMP8m were all sufficiently bright for use with systemic viruses (**Fig. 2a, Supplementary Video 1**). The expression of these indicators was visible two weeks after injection, and the expression remained stable across weeks (**Fig. 1C**), with no indication of intracellular aggregation. Of these, jGCaMP7s displayed the lowest neuropil signal but also the slowest response kinetics, as reported earlier^2^. While jGCaMP8f is reported to be brighter than previous “fast” iterations, it was not sufficiently bright for imaging. Similarly to GCaMP6f, we attempted to inject a higher volume spread out over several days, but this did not improve the brightness sufficiently. Importantly, post-mortem histological analysis indicated that brightness of the GECI was the determining factor, as the expression of GCaMP when labeled with a GFP antibody was comparable between GECIs with very different *in vivo* performance (**Fig. 2b**). Overall, the histology and *in vivo* imaging matched previous reports on PHP.eB infection^19^, with fairly even expression across the cortex and the highest density in cortical layer 5 (**Fig. 1d, 2b**). This was true for the visual cortex, somatosensory, retrosplenial and motor cortex. (**Fig. S2**). In the hippocampus, we observed almost no labeling, apart from very dense labeling in area CA2 (**Fig. S2**).

Previous reports on transgenic GECI expressing mouse lines have shown that ictal events can occur frequently in such models. In contrast to these models, none of the GECIs we tested by RO injection showed signs of ictal activity (measured by widefield fluorescence imaging) (example data shown in **Fig S1**).

### Soma targeting of existing GECIs

While sufficient brightness for imaging is a requirement for any GECI to be viable, there are many factors to consider when selecting the optimal sensor for a given experiment. The newer iterations of jGCaMP feature improved kinetics and a much higher ΔF/F relative to past versions. Accordingly, provided that brightness is sufficient, the most recent jGCaMP iteration, jGCaMP8, is preferred over past versions. However, we also observed a substantial amount of background signal, which is usually attributed to localization of the GECI to neuronal processes (neuropil). To combat this issue, we first tested three recently developed soma-targeted GCaMPs; Ribo-GCaMP6m^17^, Soma-GCaMP6f and Soma-GCaMP7f that confine GCaMP to the soma to effectively reduce background noise from neuropil^16^. We observed reduced neuropil signals in locally injected animals, but none of these were sufficiently bright for in-vivo imaging following RO AAV injection (**Table 1**).

### Expression of novel soma-targeted GECIs

We hypothesized that the brightest GECIs, the new GCaMP8s, would also work in combination with the EE-RR Soma tag. We therefore constructed Soma (EE-RR) tagged versions of the most promising jGCaMP8 variants (jGCaMP8m and s). The EE-RR Soma-jGCaMP8 was comparably bright to the unaltered jGCaMP8 at 2-4 weeks post injection, with an improved signal to noise ratio (**Fig. 3, left and middle panel, Supplementary Video S2 and 3**). In addition to improving the distinction between individual neurons, the reduced neuropil signal allowed us to image dense populations of neurons at greater depths without increasing laser power (**Fig 3, lower panel**).

**Figure 3:**
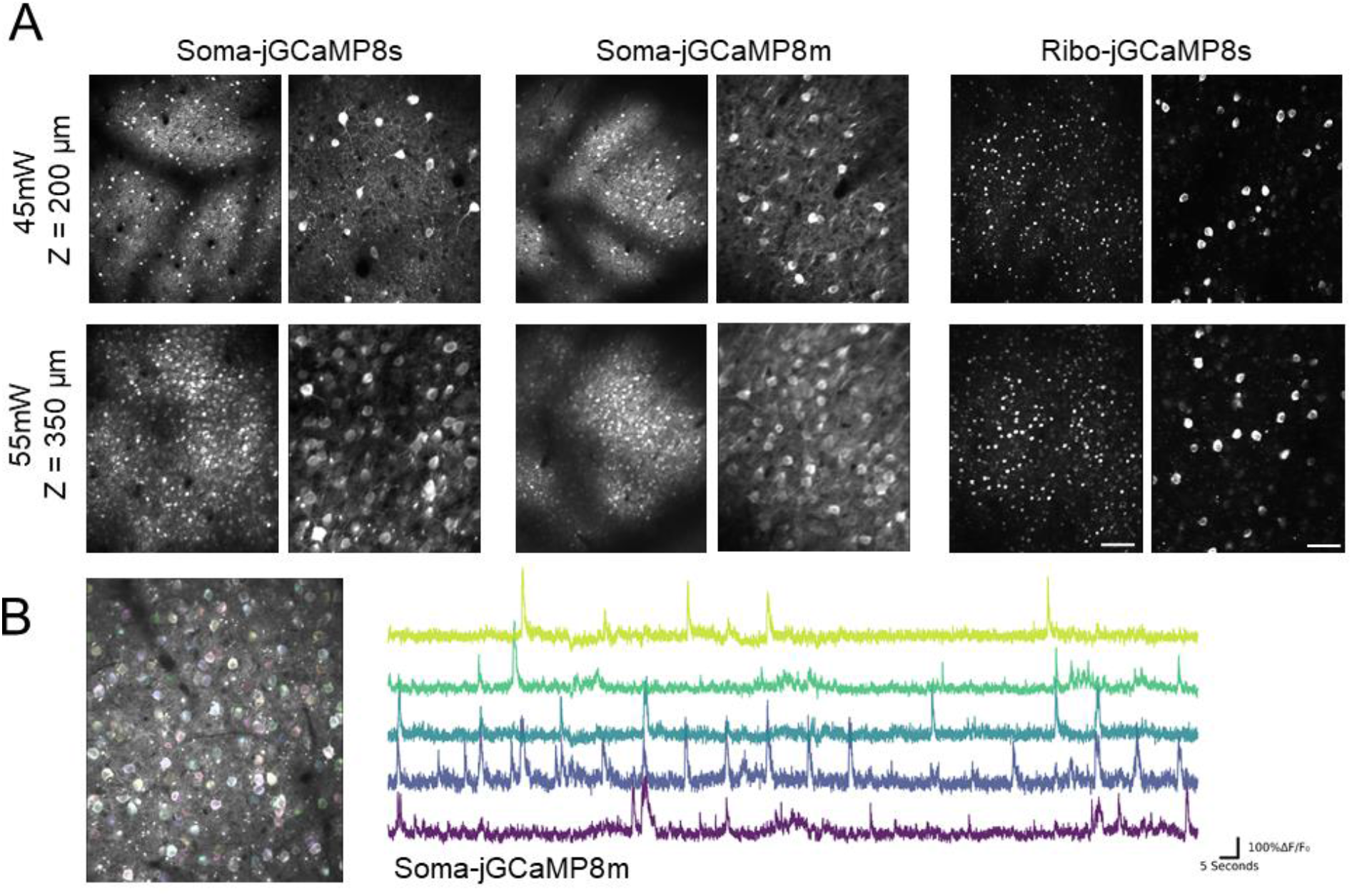
Soma-targeted GECIs expressed by retro orbital injections. **A**: Left and middle panels show EE-RR soma targeting at two imaging depths and magnifications, while the right panel shows ribosome-tethered expression. For both methods we observed high signal-to-noise and strong expression after six weeks. At 350μm below the surface the density of labeled cells was very high. In contrast to the EE-RR soma-targeted approach, ribosome-tethered GCaMP was not detectable until 4-6 weeks after injection. **B:** ROIs detected by Suite2p on the left, with five example calcium traces plotted on the right. All images shown are average intensity projections from 2000 frames with identical adjustments to brightness and contrast. Scale bars indicate 150 and 50 μm, respectively.

In comparison to EE-RR soma targeting and other non-soma targeted GECIs, ribosome-tethered GCaMP expression has been reported to show drastically reduced brightness of the attached GCaMP, demanding high laser intensity for *in vivo* imaging. However, the reported specificity of Ribo-GCaMP expression is more restricted to the soma, making it more favorable for population imaging. We therefore constructed Ribo-jGCaMP8m and s (RL10) and tested their suitability for systemic injections. In line with the previously observed reduction of brightness of GCaMP6m by the Ribo tag, ribo-tagged GCaMPs only displayed dim signal confined to a small space in the soma 2 weeks after injection. After 4-6 weeks the signal had improved, and was sufficiently bright at reasonable laser power, but the expression was still sparse relative to the EE-RR soma targeting. As previously reported for local injections of Ribo-GCaMP6m, the expression was strictly confined to the soma, with little to no visible neuropil signal (**Fig. 3, right panel, Fig S3**). However, we observed indications of aggregated GCaMP, possibly from projecting axon terminals. The widefield signals from mice injected systemically with Ribo-jGCaMPs were very weak throughout the experiment (**example image from Ribo-jGCaMP8s shown in Fig S1A**), indicating that widefield Ca2^+^ imaging using these sensors would require far more sensitive imaging equipment compared to other versions of jGCaMP8.

Ribo-jGCaMP8f and Soma-jGCaMP8f plasmids were also created, but were not tested in this paper due to the low brightness of jGCaMP8f when tested with RO injections.

### Local injections of novel soma-targeted GECIs

Despite the promising nature of RO injected GECIs for population imaging, some experiments still require local expression for Ca^2+^ imaging. We therefore tested our adapted constructs for EE-RR and ribosome-tethered jGCaMP8 and compared their performance with regular jGCaMP8 after local virus injections. Similar to the RO injected animals, we observed strong signals from both EE-RR and regular jGCaMP8 (**Fig. 4, Supplementary Video S4**), with reduced neuropil in the EE-RR version. Ribo-jGCaMP8 was relatively dim 2 and 4 weeks after injection, but after 6 weeks showed high brightness. This remained remarkably stable across time, up to the last sampling point 10 weeks after the injection. Moreover, both Ribo-jGCaMP8 versions were highly selective for expression limited to the soma, with no visible neuropil signal (**Fig. 4, Fig. S3, Supplementary Video S5 and S6**). Notably, in contrast to previous reports and our own data on Ribo-GCaMP6m, both Ribo-jGCaMP8 versions showed sufficient brightness after 6 weeks for imaging to be performed using relatively low laser power (35-45 mW).

**Figure 4:**
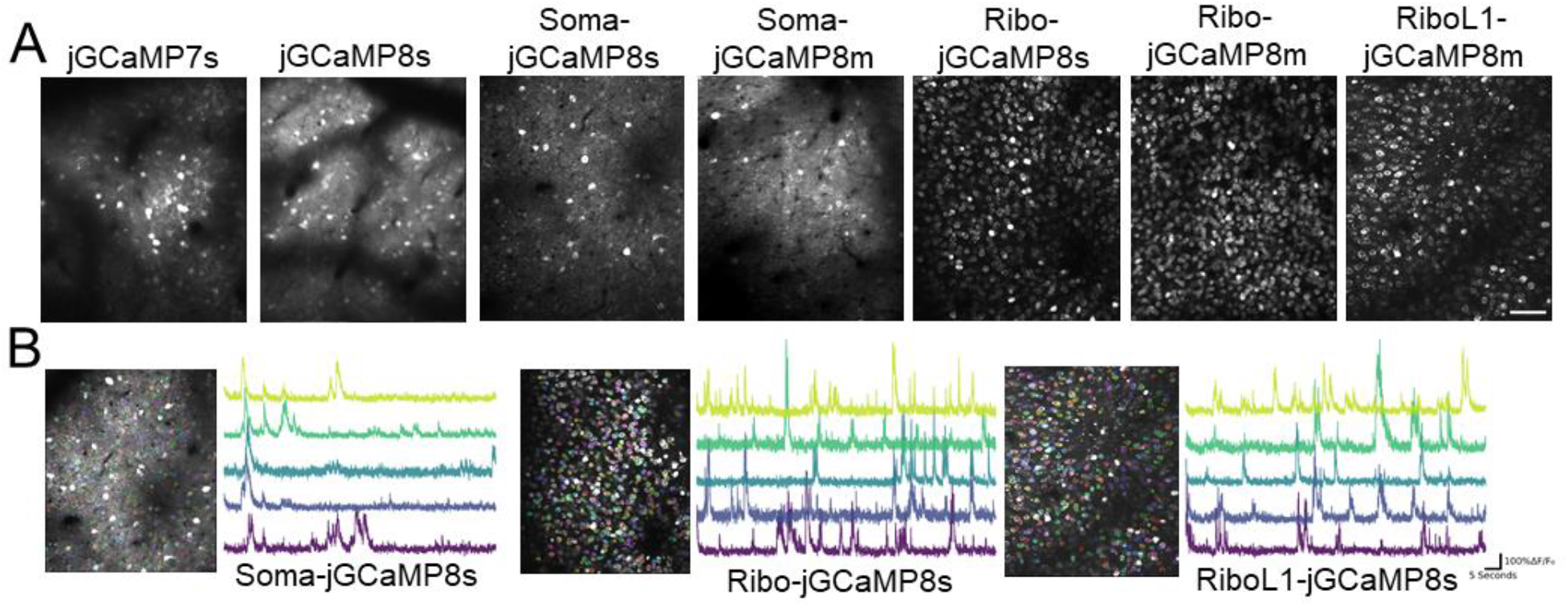
GECIs expressed by local injection of PHP.eb virus. **A:** Example images from upper layers of primary visual cortex acquired six weeks after virus injection. jGCaMP7s and 8s (left panels) were very bright but showed strong neuropil contamination of the signal. EE-RR soma targeting (middle panels) improved this somewhat, while ribosome-tethering led to a dramatic improvement. All images shown are average intensity projections from 2000 frames with identical adjustments to brightness and contrast. Scale bar indicates 75 μm. **B:** Examples of automatic cell detection from Suite2p and example traces of Ca2+ activity.

The slow expression of Ribo-jGCaMP8 may be a limiting factor to some experiments, for example by preventing imaging experiments in young animals, requiring removal of bone growth in suboptimal cranial window implants or having to perform the virus injections and window implant in separate surgeries. In an attempt to improve this we replaced the linker region of Ribo-GCaMP8 with three different linker sequences; one longer and more flexible sequence, and two variations on rigid, helical linkers. The rigid helical linkers failed to rescue Ribo-GCaMP8 expression (data not shown). However, the more flexible and longer linker, identical in amino acid sequence to the one used in EE-RR Soma-GCaMP, greatly increased the rate of protein expression. This new construct, which we term RiboL1-jGCaMP8s, showed strong expression just one week after local virus injection (**Fig. 4, Supplementary Video S7**). The expression remained stable across many weeks similar to Ribo-jGCaMP8. However, when tested with systemic injections, RiboL1-jGCaMP8 was relatively dim and the expression sparse, similar to the original Ribo-jGCaMP8 construct (data not shown).

### High signal-to-noise using novel soma-targeted GECIs

We next quantified Δf/f0 from the neuropil surrounding each neuron and compared it to the somatic signal for a selection of the GECIs tested. The boundaries used were defined by Suite2p. In line with our early observations, we found that both Ribo- and soma-targeted GECIs had higher signal-to-noise compared to jGCaMP8s (**Fig. 5**).

**Figure 5:**
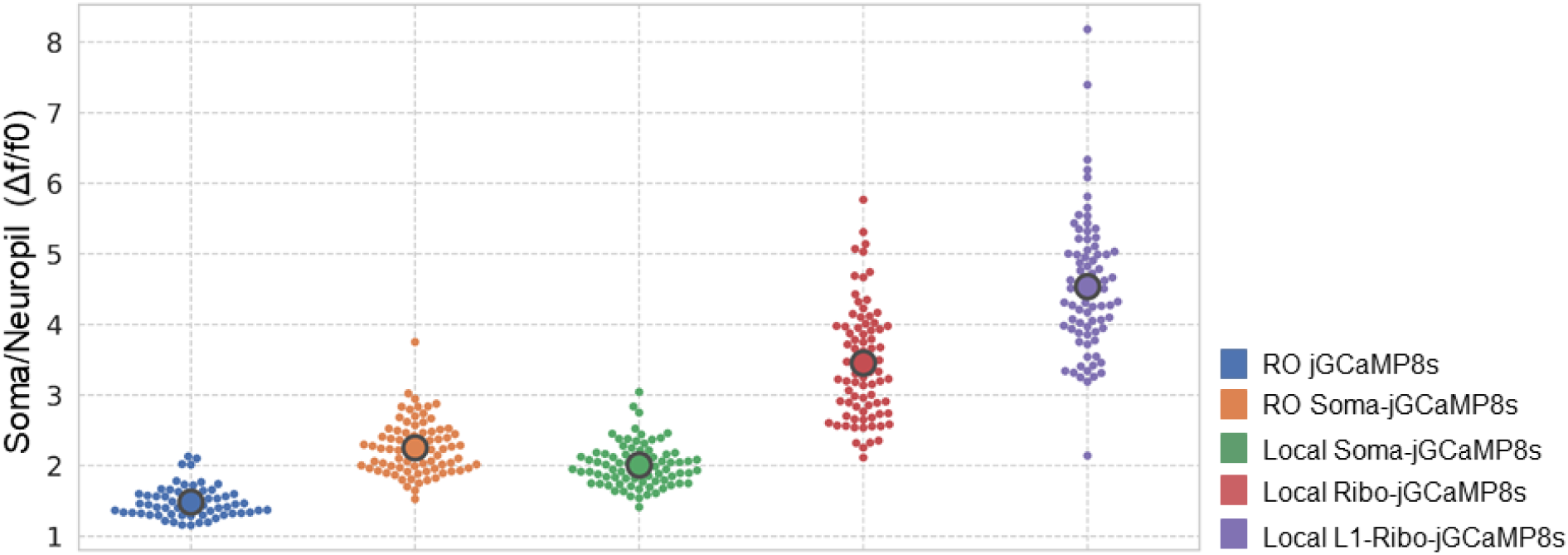
Quantification of signal-to-noise in GECIs. ΔF/F0 in soma over ΔF/F0 in neuropil, where f0 is defined as the lower fifteenth percentile of Fsoma or Fneuropil respectively. Data from 60-80 cells from single animals for each sensor.

### Applications of systemic GECI injections

One of the challenges with using traditional approaches to express GECIs is combining several transgenic constructs; local injections of two or more viruses often leads to competition and low co-expression, while transgenic GECI animals prevent the use of other transgenic lines due to the driver lines required for uniform GECI expression. To test the suitability of RO injections for this purpose, we used PV-Cre mice that express Cre under the parvalbumin (PV) promoter, and performed an RO injection of Soma-jGCaMP8 combined with local injections of an AAV5 vector expressing a flexed hM4D DREADD receptor. Indeed, this led to co-expression of both constructs in putative PV neurons and uniform expression of jGCaMP8 in surrounding neurons (**Fig. 6a**). This was verified by post-mortem histology (**Fig 6b**).

**Figure 6:**
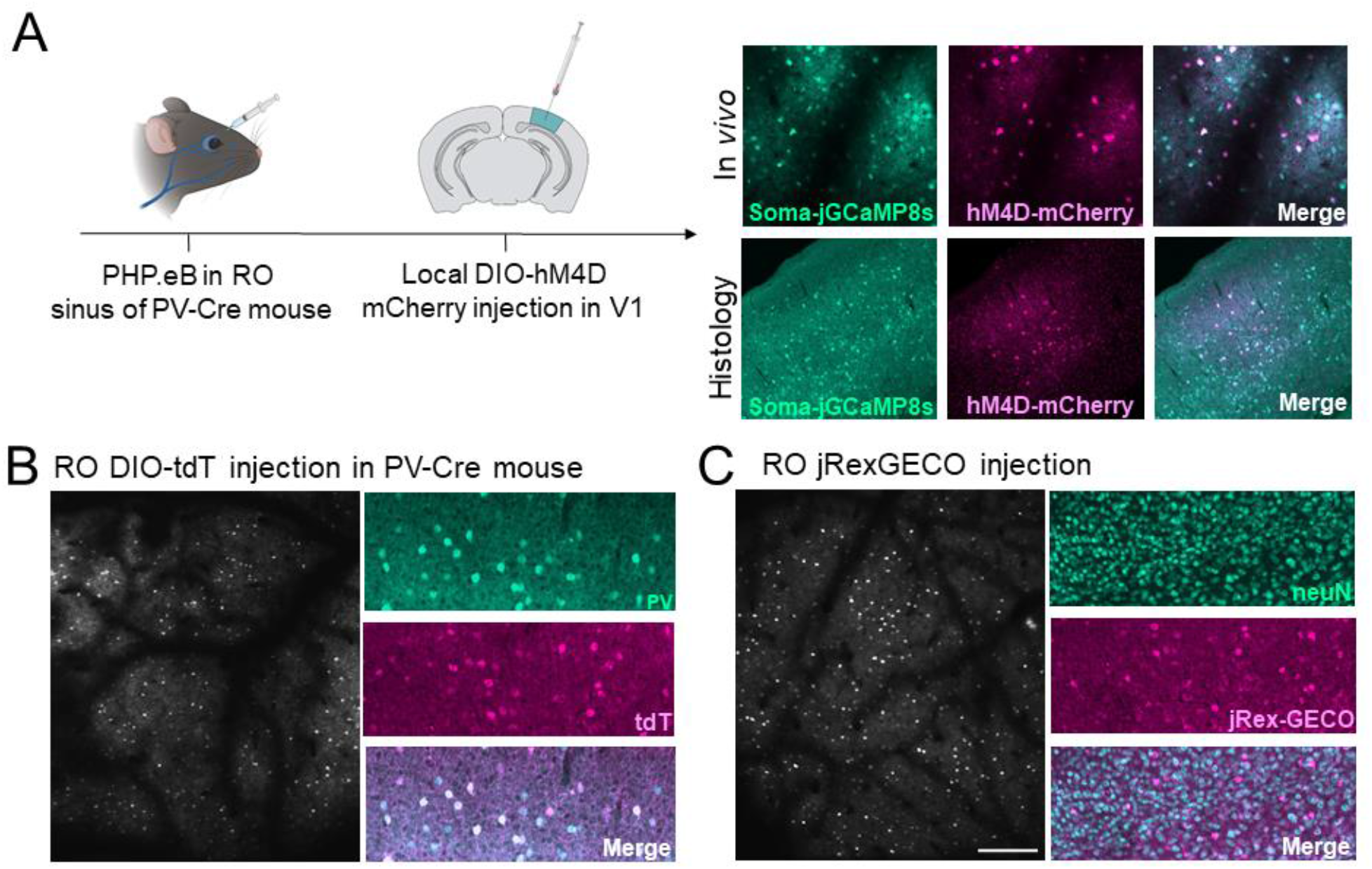
Applications of systemic GECI expression. **A:** experimental overview for GECIs expressed by RO injection and Cre-dependent hM4D expressed by local injection. **B:** RO injection of Cre-dependent TdTomato in PV-Cre mice led to strong expression of tdTomato across cortex (left: example average intensity projection of 2000 frames). The selectivity to PV expressing neurons was verified by histology (right panel). **C:** RO injection of the red-shifted GECI jREX-GECO1. Scale bar indicates 250 μm.

Another opportunity by combining uniform GECI expression with other transgenics is to identify specific populations of neurons by labeling with a fluorescent protein outside the color range of traditional GCaMPs. To this end, we again used PV-Cre mice and performed RO injections of a Cre-dependent TdTomato PHP.eB virus. We observed highly selective labeling of PV neurons, in both *in vivo* and histological samples (**Fig. 6c**). Next, we tested the red-shifted GECI jREX-GECO1, a long stokes-shift version of the red GECI RGECO that is optimized for two-color imaging with a single laser source. Using the same 920nm laser as for green GECIs, jREX-GECO1 was even brighter than jGCaMP8s. This indicates that jREX-GECO1 could be a viable GECI to use in combination with imaging axonal activity using a green-shifted Ca^2+^ indicator (e.g.^20,21^).

Finally, the reduced neuropil expression of our soma-targeted constructs opens for detailed investigations of input-output relationships in neurons by combining these GECIs with axon-targeted^20^ expression of a red-shifted GECI. To this end, we constructed axon targeted jREX-GECO1 (hSyn-Axon-jREX-GECO1) and performed local virus injections into the dorsal lateral geniculate nucleus (dLGN), combined with local injections of RiboL1-jGCaMP8s in V1. We then imaged the activity in axonal boutons and cell somas in V1 two weeks after viral injections, and found strong signals from both GECIs (**Fig 7a and b** shows examples from *in vivo* imaging and histology, respectively). We detected single axonal boutons with both highly correlated and non-correlated changes in fluorescence with the soma signal (**Fig 7a and c**). This indicates that our soma-targeted GECIs could be viable for use in combination with imaging axonal activity using e.g. Axon-jREX-GECO1. Axon-jGCaMP8s and m were also created and will be available on Addgene, but have not been tested in this paper.

**Figure 7:**
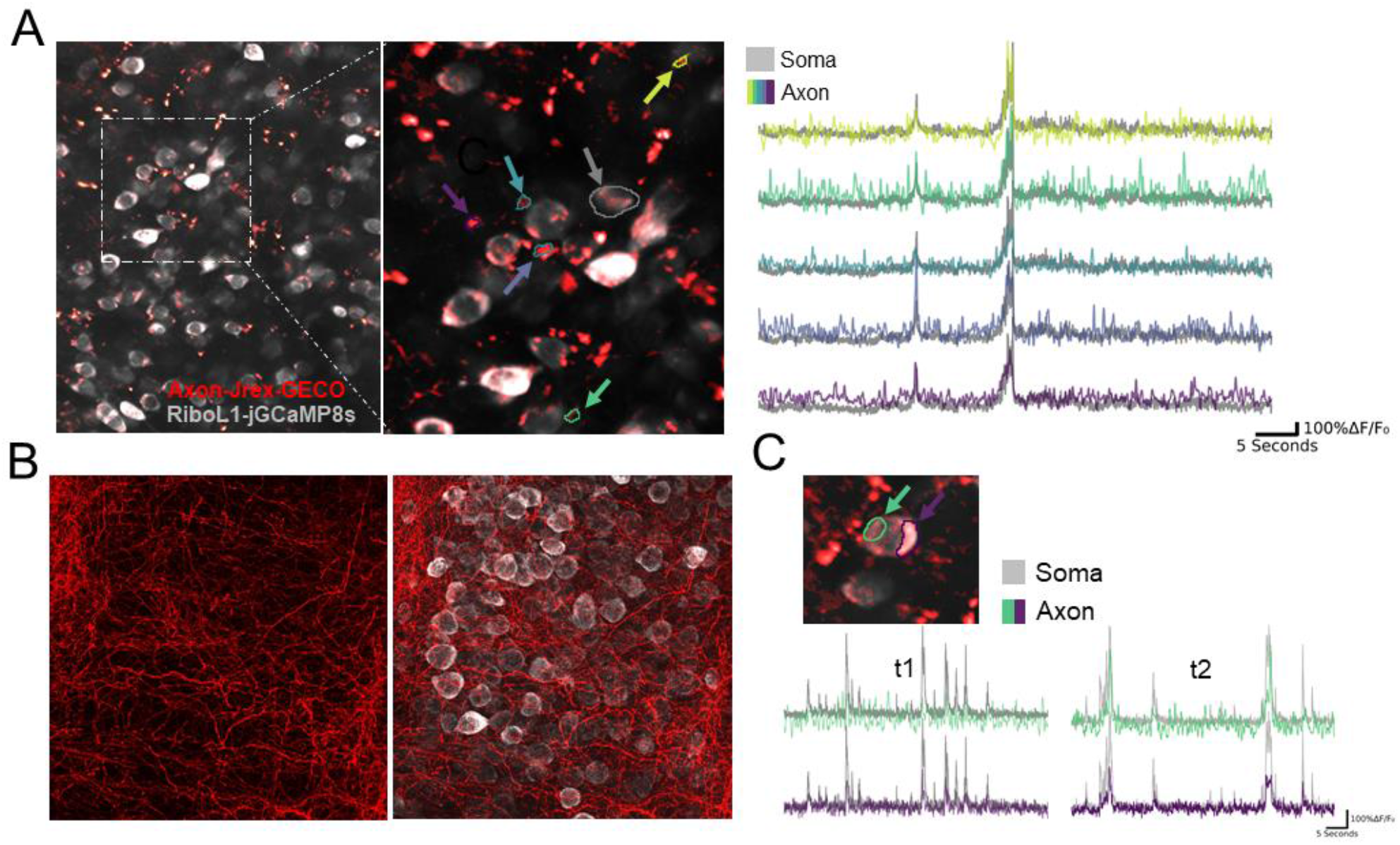
Simultaneous in *vivo* imaging of axonal and somatic activity. **A) Left panel:** An example field-of-view in primary visual cortex with Axon-jREX-GECO1 in red and RiboL1-jGCaMP8s in grey. Colored arrows indicate labeled regions of interest from the red channel. The cell soma ROI from the green channel is indicated by a grey arrow. **Right panel:** Red channel Ca^2+^ traces (colored) from each bouton overlaid on the soma signal (grey). Despite clear spatial separation, the indicated regions of interest show similar activity patterns (middle and right panel). **B:** Histological verification of expression. **C:** Two regions of interest in close proximity to a cell soma, indicated by colored arrows, and corresponding Ca^2+^ traces overlaid on the soma signal at different time points.

## DISCUSSION

Engineered AAV serotypes with high affinity for the central nervous system that can be delivered intravenously represent a minimally invasive and low-cost method for introducing genetic payloads into the brain. Yet, these new serotypes, notably PHP.eB, have not been much used to deliver GECIs, despite obvious advantages in terms of animal welfare, cost, productivity and experimental flexibility. Previous work shows that widefield imaging with systemically administered GCaMP6f is feasible using other promoters than synapsin^22,14^. In our own preliminary experiments we experienced that typical GECIs (such as GCamp6f) were not bright enough to be compatible with systemic administration. Here, we screened 14 GECIs and 2 fluorescent probes and show that newer iterations of jGCaMPs, particularly jGCaMP7s, jGCaMP8s and m, are indeed sufficiently bright for two-photon in-vivo calcium imaging when administered intravenously in PHP.eB AAVs. We also observed strong neuropil signals from these sensors which may influence accuracy of individual neuron activity, regardless of administration route. By fusing the latest jGCaMP variants with soma-targeting peptides we overcame this issue. Remarkably, we show that our Soma-jGCaMP8 and Ribo-jGCaMP8 outperform existing GECIs using both systemic and local virus injections.

An intravenous AAV injection in the retro-orbital sinus can be performed in a minute or two and requires very little training. The procedure is substantially less invasive than stereotaxic/intracranial injections. For the work presented here, one person successfully performed RO injections in 16 animals in 45 minutes. The intravenous injection provides largely uniform expression across the mouse brain, which is stable over extended time periods. This stands in contrast to local virus injections that often result in excessive expression leading to cell death. We note that in our hands, stable expression was resulting both from the administration route and the PHP.eB serotype. The expression of locally injected PHP.eB GECIs is also far more shielded from over-expression and toxicity compared to traditional AAV serotypes like AAV9, albeit not to the same level as RO injected PHP.eB virus.

Despite the many advantages of intravenously injected virus for GECI delivery, it has one considerable drawback; intravenous administration requires a large dose of virus per animal. If all viruses are purchased from commercial vendors, this could be prohibitively expensive. On the other hand, if viruses are produced in-house or by a local virus core, scaling up production to suitable levels is relatively inexpensive, and was not an issue for our experiments. Performing injections in younger animals will substantially reduce the amount of virus required. Another strategy which has been employed, is to use a stronger promoter such as CAG^23^. Unlike the Synapsin promoter which is commonly used in neuroscience, CAG would not limit expression to neurons, and despite the strong tropism of PHP.eB AAV to neurons, this would result in some glial cells expressing GCaMP. This could be prevented if a flexed construct is used, but this would introduce the need for transgenic lines or an additional Cre expressing virus. An additional caveat concerning the PHP.eB and AAV9 is that we find a clear bias in expression for cortical layer 5, striatum, CA2 and subiculum regions. While the bias towards cortical layer 5 might be explained by the large cell volumes and thus higher capacity for transgene production, this does not appear to be a common feature for the preferred brain areas. An alternative explanation might be that differences in vascularization is determinant for expression levels. However, this is at least unlikely for CA2 of the hippocampus which exhibits strikingly strong expression compared to neighboring hippocampal areas with no large differences in vascularization^24,25^. Notably, the bias for layer 5 neurons was not as clear for ribosome-tethered versions of jGCaMP8. Because transfection efficiency is mainly decided by serotype, this could indicate that it could be a result of signal masking by neuropil. If the reason for expression differences are identified and reduced in future iterations of synthetic AAV serotypes, less virus may be required to achieve sufficient brightness in the upper layers of cortex for *in vivo* imaging.

The need for high brightness of the GECI for systemic administration also caused neuropil contamination of the signals. We therefore made use of two soma-targeting strategies that restricted expression to the cell soma. Moreover, we show that EE-RR soma-targeting led to highly stable expression which was already visible after two weeks, while ribosome-tethering reduced the brightness to such an extent that we did not observe cells during in-vivo imaging until 4-6 weeks after injection. Nonetheless, imaging could be performed using reasonable laser power 6 weeks after injection, and the ribosome-tethering led to highly selective although somewhat limited expression in the cell soma. For local injections, ribosome-tethering showed very high density of cell labeling, but again the expression was slow. We therefore introduced a novel version with a modified linker region (RiboL1-jGCaMP8) that greatly improved this feature. Using RiboL1-jGCaMP8 we could initiate imaging just one week after local virus injection, with no apparent drawbacks such as over-expression over extended time periods. In general, soma-targeting GECI expression leads to improved signal-to-noise and the possibility to detect activity from a higher number of cells as their activity is not masked by neuropil activity. We also observed that automatic cell detection in suite2p was more accurate and required smaller data sets.

A major challenge when using transgenic animal models to express GECIs is the need for driver lines with general promoters preventing the use of other transgenics. Moreover, co-expression of several viruses, at least in our hands, often proves difficult. In contrast, we show that RO injected PHP.eB virus is compatible with transgenic lines and co-expression of another virus to obtain cell specific expression of e.g. a chemogenetic receptor to manipulate their activity or labelling a specific cell population.

In summary, we present a suite of viral vectors for use with both systemic and local administration that show remarkably high performance and sustainable expression over longer periods. Due to the simplicity of the methods, high experimental flexibility, low cost and high performance, we believe that these soma-targeted GECI constructs are promising candidates to replace transgenic animal models for GECI expression.

## MATERIALS & METHODS

### GECI Plasmids

All plasmids were transformed into NEB Stable (NEB) competent cells for amplification, and purified using the Zymopure II maxiprep kit (Zymo Research). To obtain “Soma” tagged GCaMP8, pAAV-Syn-SomaGCaMP7 was digested with HpaI and EcoRI to isolate the linker and Soma-tag. The fragment was then ligated into AAV-hSyn-GCaMP8s, m and f, which was previously digested using the same restriction enzymes. pAAV-Syn-SomaGCaMP7^16^ was a gift from Edward Boyden (http://n2t.net/addgene:158759;RRID:Addgene_158759). AAV-syn-jGCaMP8s-WPRE^18^ was a gift from GENIE Project (http://n2t.net/addgene:162374; RRID:Addgene_162374), as well as jGCaMP8f (Addgene:162376) and jGCaMP8m (Addgene:162376). To obtain “Ribo” tagged GCaMP8, pyc126m (Ribo-GCaMP6m) was digested with HpaI and EcoRI to isolate the linker and Ribo-tag. The fragment was then ligated into AAV-syn-GCaMP8s, m and f, previously linearized using the same restriction enzymes. pycm126^17^ was a gift from Jennifer Garrison & Zachary Knight (http://n2t.net/addgene:158777;RRID:Addgene_158777). To obtain Synapsin promoter expressed jREX-GECO1, the jREX-GECO1 coding sequence was cut from CMV-jREX-GECO1 using BamHI and EcoRI, and inserted into a pAAV-Syn-Chr2 plasmid, which was digested with the same restriction enzymes, removing the coding sequence of Chr2 and replacing it with jREX-GECO1. jREX-GECO1 expressed under a CMV promoter was a gift from Neurophotonics^26,27^. The hSyn plasmid, pAAV-Syn_ChR2(H134R)-GFP^28^ was a gift from Edward Boyden (http://n2t.net/addgene:58880;RRID:Addgene_58880). To obtain Ribo-GCaMP8 with modified linkers, three different linkers were synthesized (Table S1, GeneArt Invitrogen, codon optimized) and inserted into GCaMP8s, m and f plasmids using HpaI and AccI (NEB). The first linker, RiboL1, was adapted from the SomaGCaMP7f plasmid. The second and third linkers were variations of rigid helical linkers^29^, RiboL2: LEA(EAAAK)4ALE, and RiboL3: LEA(EAAAK)4ALEA(EAAAK)4ALE (data not shown for RiboL2 and L3). To obtain axon targeted jREX-GECO1 and jGCaMP8, a fragment (Table S1) encoding the 20AA axon targeting motif from Broussard et al.,^20^ was synthesized (Invitrogen, GeneArt) and ligated into jREX-GECO1 and GCaMP8s and m plasmids. Additional plasmids, GCaMP6f, somaGCaMP6f, jGCaMP7f, jGCaMP7s, mNeonGreen and flex-tdTomato were acquired from addgene and were not modified in this paper. pAAV.Syn.GCaMP6f.WPRE.SV40^30^ was a gift from Douglas Kim & GENIE Project (http://n2t.net/addgene:100837;RRID:Addgene_100837). pGP-AAV-Syn-jGCaMP7f-WPRE was a gift from Douglas Kim & GENIE Project (http://n2t.net/addgene:104488; RRID:Addgene_104488). pGP-AAV-Syn-jGCaMP7s-WPRE^2^ was a gift from Douglas Kim & GENIE Project (http://n2t.net/addgene:104487; RRID:Addgene_104487). pAAV-CAG-mNeonGreen^13^ was a gift from Viviana Gradinaru (http://n2t.net/addgene:99134; RRID:Addgene_99134). pAAV-FLEX-tdTomato was a gift from Edward Boyden (http://n2t.net/addgene:28306; RRID:Addgene_28306). Plasmids for AAV packaging were acquired from Addgene and Penn Vector core, which is now available from Addgene. Only PHP.eB serotype viruses were used in this paper, except for the cre-dependent DREADD-mCherry, pAAV-hSyn-DIO-hM4D(Gi)-mCherry^31^ which was a gift from Bryan Roth (Addgene viral prep # 44362-AAV5;http://n2t.net/addgene:44362;RRID:Addgene_44362). The PHP.eB serotype plasmid, pUCmini-iCAP-PHP.eB^13^ was a gift from Viviana Gradinaru (http://n2t.net/addgene:103005;RRID:Addgene_103005). The DeltaF6 helper plasmid, pAdDeltaF6, was a gift from James M. Wilson (http://n2t.net/addgene:112867;RRID:Addgene_112867).

#### AAV production

Viral vectors were produced in house according to the protocol developed by Rosemary C Challis et al^32^. Briefly, AAV HEK293T cells (Agilent) were cultured in DMEM with 4.5 g/L glucose & L-Glutamine (Lonza), 10% FBS (Sigma) and 1% PenStrep (Sigma), in a 37°C humidified incubator. The cells were thawed fresh and split at ∼80% confluency until four 182.5cm^2^ flasks were obtained for each viral prep. The cells were transfected at 80% confluency and the media was exchanged for fresh media directly before transfection. The cells were triple transfected with dF6 helper plasmid and PHP.eB serotype plasmid. Polyethylenimine (PEI), linear, molecular weight (MW) 25,000 (Polysciences, cat. no. 23966-1) was used as the transfection reagent. Media was harvested three days after transfection and kept at 4°C, and media with cells was harvested five days after transfection, and combined with the first media harvest. After 30 min centrifugation at 4000 g, the cell pellet was incubated with SAN enzyme (Arctic enzymes) for 1 hour. The supernatant was mixed 1:5 with PEG and incubated for 2 hours on ice, then centrifuged at 4000g for 30 minutes to obtain a PEG pellet containing the virus. The PEG pellet was dissolved in SAN buffer and combined with the SAN cell pellet for incubation at 37°C for 30 minutes. To purify the AAV particles, the suspension was centrifuged at 3000 g for 15 minutes. The supernatant was loaded on the top layer of an Optiseal tube with gradients consisting of 15%, 25%, 40% and 60% iodixanol (Optiprep). Ultracentrifugation was performed for 2.5 hours in 18°C at 350’000g in a type 70 Ti rotor. The interface between the 60% and 40% gradient was extracted along with the 40% layer, avoiding the protein layer on top of the 40% layer. The viral solution was filtered through a Millex-33mm PES filter before adding to an Amicon Ultra-15 centrifugal filter device (100-kDa molecular weight cutoff, Millipore). A total of four washes with 13 ml DPBS were performed at 3000 g before concentration to a volume of ∼750ul. Viral solutions were sterilized through 13 mm PES syringe filters 0.2 μm (Nalgene), and stored in sterile screw-cap vials at 4°C.

Viral titres were determined using qPCR with primers targeting AAV2 ITR sites^33^ (Table S1), following the <u>“AAV Titration by qPCR Using SYBR Green Technology”</u> protocol by Addgene^10^. Briefly, 5 ul of viral sample was added to 39 ul ultrapure H2O, 5 ul 10x DNase buffer, 1 ul DNase, and incubated at 37°C for 30 minutes to eliminate all DNA not packaged into AAV capsids. 5 ul of the DNase treated sample was added to a reaction mix consisting of 10 ul 2x SYBR master mix, 0.15 ul of each primer (100uM) and 4.7 ul nuclease free H2O. Cycling conditions for the qPCR program were: 98°C 3 min / 98°C 15 sec / 58°C 30 sec / read plate/ repeat 39x from step 3 / melt curve.

In addition to the constructs tested in the manuscript, we also made Soma-jGCaMP8f and Ribo-jGCaMP8f. All plasmids will be deposited to Addgene.

#### Experimental animals

All work with experimental animals was performed at the animal facility at the Department of Biosciences, Oslo, Norway, in agreement with guidelines for work with laboratory animals described by the European union (directive 2010/63/EU) and the Norwegian Animal Welfare Act from 2010. The experiments were approved by the National Animal Research Authority of Norway (Mattilsynet, FOTS ID 14680).

Four weeks old male C57/BL6j mice were purchased from Janvier Labs, and housed in GM500 IVC cages in groups of four. After an acclimation period of two weeks, the animals were split into two mice per cage prior to virus injections. One week after injections, the mice were housed individually, and remained single-housed for the duration of the experimental period. The housing room had a 12/12 hour light cycle, with lights off at noon. In the light phase, light intensity in the room was 215 lux, and in the cages varied from 20-60 lux, depending on position in the rack. All experiments were performed in the dark phase. For enrichment purposes, each cage had a running wheel and large amounts of nesting material, and the mice had *ad libitum* access to food and water.

#### Retro-orbital injections

Pairs of mice were randomly assigned to a viral vector. The mice were placed in an induction chamber and briefly anesthetized by isoflurane, before they were transferred to a mask with 1-2% isoflurane delivered. An eye drop of local anesthetic (oxybuprocaine 4mg/mL, Bausch Health), was applied to the right eye, and one minute later 100-150 μL of virus injected into the retro-orbital sinus using a U100 insulin syringe (BD micro-fine 0.3mL, 30 gauge needle). The volume was determined based on the animal’s weight^15^. The surface of the eye was flushed with saline and cleaned with a cotton tip. The mice were then placed back in the home cage and monitored for 10-15 minutes, before they were returned to the housing room. All animals fully recovered within minutes. In one single mouse, we observed eye damage to the injected side after one week. It is not clear whether this resulted from the injection or resulting from the high incidence of eye abnormalities in the c57bl6 mouse strain^34^. Each viral vector was tested in at least two mice with RO injections and one mouse with local injection.

#### Surgical procedures

The mice were anesthetized by an intraperitoneal injection of a ketamine/xylazine cocktail (Ketamine 12.5 mg/kg, Pfizer; xylazine 5mg/kg, Bayer Animal Health GmbH). The top of the head was shaved and the animals placed on a heating pad in a stereotaxic frame with a mouse adapter (Model 926, David Kopf Instruments). The eyes were covered with white vaseline to prevent drying and to protect them from light. Dexamethasone (5 mg/kg, MSD Animal Health) was delivered via an intramuscular injection to prevent edema, and local anesthetic bupivacaine (Aspen) injected in the scalp. In a subset of animals, the mice were anesthetized by isoflurane (3.5% induction, 1-1.5% maintenance) and additionally injected subcutaneously with buprenorphine (0.05mg/kg, Indivior Ltd) for analgesia. The skin was cleaned with 70% ethanol and chlorhexidine, and a small piece of skin covering the top of the skull was cut away. The periosteum and other membranes were removed using fine forceps and cotton swabs, and the surface of the skull slightly scored with a scalpel. A custom titanium head post was attached using a few drops of cyanoacrylate, and secured using VetBond (3M) and C&B Metabond (Parkell). A 3.0 mm craniotomy was made using a Perfecta 300 hand-held drill (W&H) with a 0.5 mm drill bit (Hager & Meisinger GmbH), centered over primary visual cortex (center coordinates were 2.5 mm ML and 1 mm AP relative to lambda). Custom cranial windows were made by attaching a 3.0 mm diameter round glass (Tower Optical) with 0.45 mm thickness to a 5.0 mm diameter glass (Warner Instruments, USA) with 100 μm thickness using Norland Optical adhesive (Thorlabs GmbH, Germany) under UV light. The cranial window was implanted and secured with C&B Metabond, and a 3D printed light shield was attached to the head post with black dental acrylic^7^. At the end of the procedures, the mice were injected subcutaneously with 0.3 mL 0.9% saline, meloxicam (5 mg/kg, Boehringer Ingelheim VetMedica GmbH) and Antisedan (0.0012 mg/kg, Orion Pharma). Injections of metacam were repeated for three days.

In a subset of mice, bone growth partially obscured the view through the cranial window over the course of the experimental period. In these cases, the animal was anesthetized by isoflurane, the window removed to clear away any bone growth and other debris, and a new cranial window implanted^7^. The procedure was performed one week prior to imaging to allow the tissue to recover from potential damage during bone removal.

#### Local virus injections

Glass capillaries (OD 1.14 mm:, ID 0.53 mm) were pulled and beveled at a 40 degree angle^35^, and mounted in a NanoJect 3 (Drummond Scientific, USA). The pipette was front loaded with the virus solution and 150 nL injected at a depth of 350-500 μm below the dura, in 5 nL steps. After the last injection, the pipette was left in the tissue for five minutes before retraction and loading of a new pipette. Two to three different constructs were injected per animal, spaced at least 700 μm apart. After the final injection, the exposed brain surface was cleaned with saline and a cranial window implanted as described above.

#### Co-expression of Axon-jREX-GECO1 and soma-targeted GECIs

For co-expression of Axon-jREX-GECO1 and RiboL1-jGCaMP8, surgical procedures were conducted as described above. After attachment of the head post, a small craniotomy was made above the dLGN, and 150nL of virus slowly injected in 5nL steps over 5 minutes (injection coordinates relative to bregma were 2.1 mm posterior, 2.3 mm lateral and 2.5 mm below the dura). A larger craniotomy was then made over V1, RiboL1-jGCaMP8s was injected and a cranial window implanted as described above.

#### Widefield imaging

Widefield imaging was used to monitor the expression levels of the Ca^2+^ sensors and quality of the cranial windows. The mice were head-fixed on a custom 3D-printed running wheel using optical posts that were mounted to the optical table, holding clamps (Standa) and modified ball-joints (Thorlabs GmbH) allowing for adjustments in AP elevation. Single images were acquired by a Canon EOS 4000D camera through a 5X Mitutoyo long working-distance objective (0.14 NA) in an Olympus BX-2 microscope. The light source was a xenon arc Lambda XL lamp (Sutter Instruments) with 480/545nm and 560/635 nm filters (#39002 and #39010, Chroma). All animals were imaged using two sets of parameters at each time point, with exposure times of 600 and 2000 ms, and ISO of 100 and 400, respectively. The mice ran freely in darkness during imaging. In addition, widefield videos were captured at 25Hz during both spontaneous activity (in darkness) and with visual stimulation.

#### Two-photon imaging

For *in vivo* two-photon imaging we used a resonant-galvo Movable Objective Microscope (MOM, Sutter Instruments) with a MaiTai DeepSee laser (SpectraPhysics) set to a wavelength of 920 nm (990 nm used for TdTomato). The main objective used for screening was a 10X objective (TL-10×2P, 0.5 NA, 7.77 mm working distance, Thorlabs), giving a field of view of approximately 1665 × 1390 μm. In mice with successful GCaMP imaging we also imaged at lower depths (200-500μm) using a Nikon 16X objective (NA 0.8), giving a field of view of approximately 1050 × 890 μm. The laser was controlled by a pockel’s cell (302 RM, Conoptics), and fluorescence detected through Chroma bandpass filters (HQ535-50-2p and HQ610-75-2p, Chroma) by PMTs (H10770PA-40, Hamamatsu). Images were acquired at 30.9 Hz using MCS software (Sutter Instruments). Output power at the front aperture of the objective was measured prior to each imaging session with a FieldMate power meter (Coherent) and set to 50 mW, unless mentioned otherwise. The microscope was tilted to an angle of 6-12 degrees during imaging to match the surface of the brain, in addition to the 4 degree forward tilt made by the head-fixing apparatus.

#### Co-expression of GCaMP and cell-specific DREADDs

PV-Cre mice (Jackson Laboratories, strain #017320) were injected with Soma-jGCaMP8s as described above. Two weeks later, 150nL of pAAV5-hSyn-DIO-hM4D(Gi)-mCherry (#44362, Addgene) was injected into the cortex, and a cranial window implanted.

#### Visual stimulus

Sinusoidal drifting gratings were generated using the open-source Python software PsychoPy^36^, and synchronized with two-photon imaging through a parallel port and PCIe-6321 data acquisition board (National Instruments). We used drifting gratings of 3 orientations (0, 135 and 270 degrees) with a spatial frequency of 4 cycles per degree and temporal frequency of 2Hz. Stimulus was shown for 3 seconds, interleaved with an 8 second grey screen period.

#### Two-photon imaging analysis

Motion-correction and automatic detection of regions of interest (ROIs) was performed using suite2p^37^. The data was then manually curated, and analyzed using custom Python scripts. ΔF/F0 was defined as (F - 0.7 * Fneu) / F0, where F is the raw fluorescence, Fneu is the neuropil fluorescence as defined by suite2p and F0 is the mean of the tenth percentile of F. The notebooks used to generate the calcium trace figures are available on Github (github.com/sverreg/calciumtrace).

#### Widefield imaging analysis

To measure changes in fluorescence over time in widefield imaging videos, we used ImageJ (Fiji). Videos were spatially downsampled, and regions of interest (ROIs) selected in the center of the cranial window. Changes in relative fluorescence was calculated by (F-F_0_/F_0_), where the baseline fluorescence (F_0_) was defined as the mean fluorescence across all frames from “spontaneous” and “visual stimulus” runs in the entire cranial window. Calcium signal traces were obtained from the average fluorescence intensity in an approximately 200μm diameter circular area.

#### Histology

Six weeks after virus injection, all animals were deeply anesthetized by an intraperitoneal injection of Euthasol (pentobarbital sodium 100 mg/kg, Le Vet) and intracardially perfused with PBS followed by 4% paraformaldehyde (PFA) in PBS. Brains were dissected out and postfixed for 24 hours followed by cryoprotection in 30% sucrose in PBS for 48 hours. 40μm coronal sections were cut with a cryostat. All sections were stained free-floating on constant agitation. The sections were rinsed three times in PBS followed by blocking in 2% bovine serum in 0.3 % Triton X-100 in PBS for 1 hour before incubation with primary antibody in blocking solution overnight (all antibodies used are listed in Table 3). After rinsing, sections were incubated with secondary antibodies in PBS for 1 hour. Sections were then mounted on Superfrost Plus adhesion slides and dried for 2 hours. After rinsing in dH_2_O and additional drying for 1 hour, sections were coverslipped with mounting medium (Ibidi). Tile scans were acquired with 20% overlap on an Andor Dragonfly spinning-disc microscope with a motorized platform, and stitched using Fusion software (Bitplane). The Andor Dragonfly was built on a Nikon TiE inverted microscope equipped with a Nikon PlanApo 10x/0.45 NA objective.

**Table 3:**
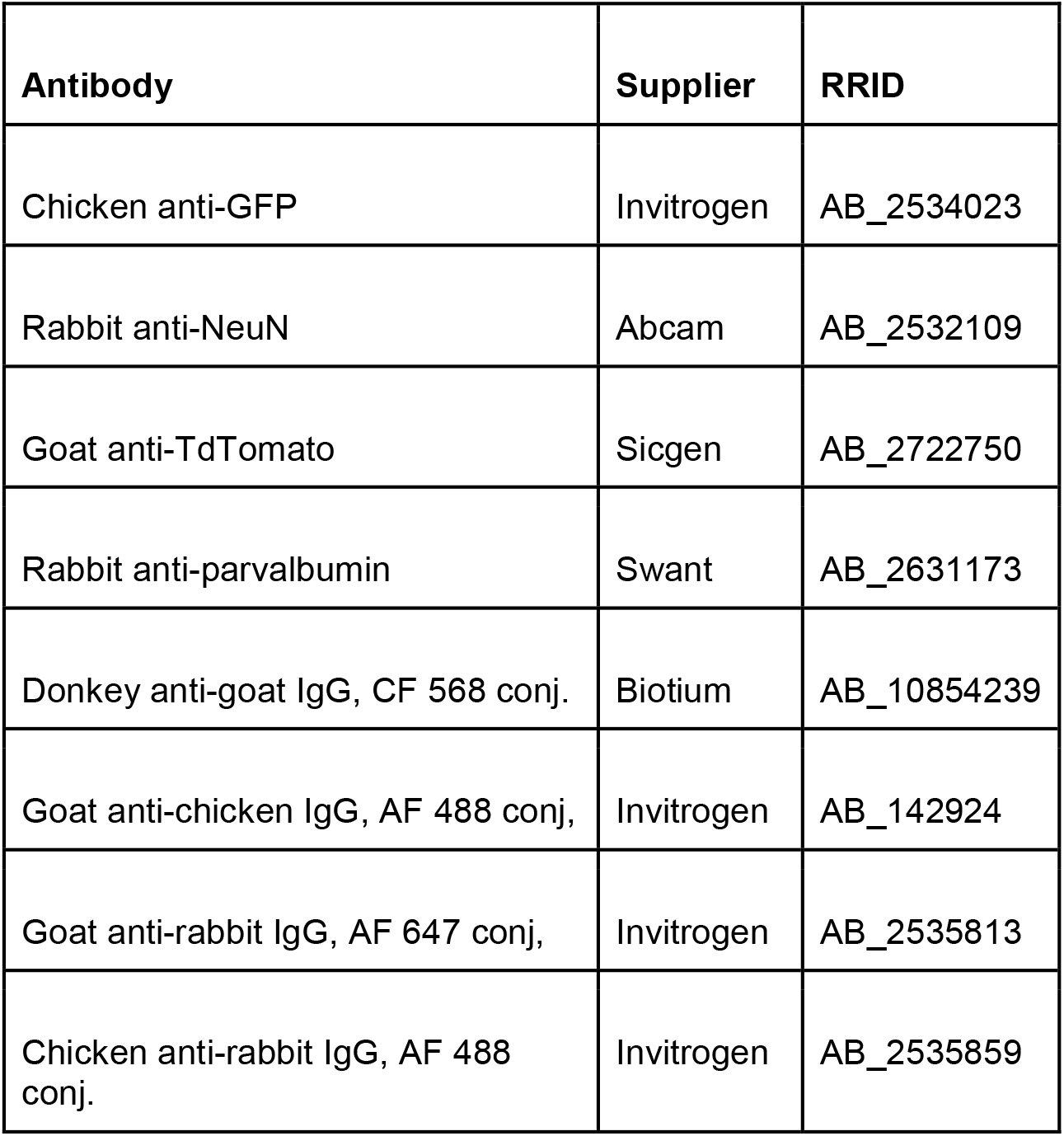
List of antibodies used for post-mortem histology.

| Antibody | Supplier | RRID |
| --- | --- | --- |
| Chicken anti-GFP | Invitrogen | AB_2534023 |
| Rabbit anti-NeuN | Abcam | AB_2532109 |
| Goat anti-TdTomato | Sicgen | AB_2722750 |
| Rabbit anti-parvalbumin | Swant | AB_2631173 |
| Donkey anti-goat IgG, CF 568 conj. | Biotium | AB_10854239 |
| Goat anti-chicken IgG, AF 488 conj, | Invitrogen | AB_142924 |
| Goat anti-rabbit IgG, AF 647 conj, | Invitrogen | AB_2535813 |
| Chicken anti-rabbit IgG, AF 488 conj. | Invitrogen | AB_2535859 |

## AUTHOR CONTRIBUTIONS

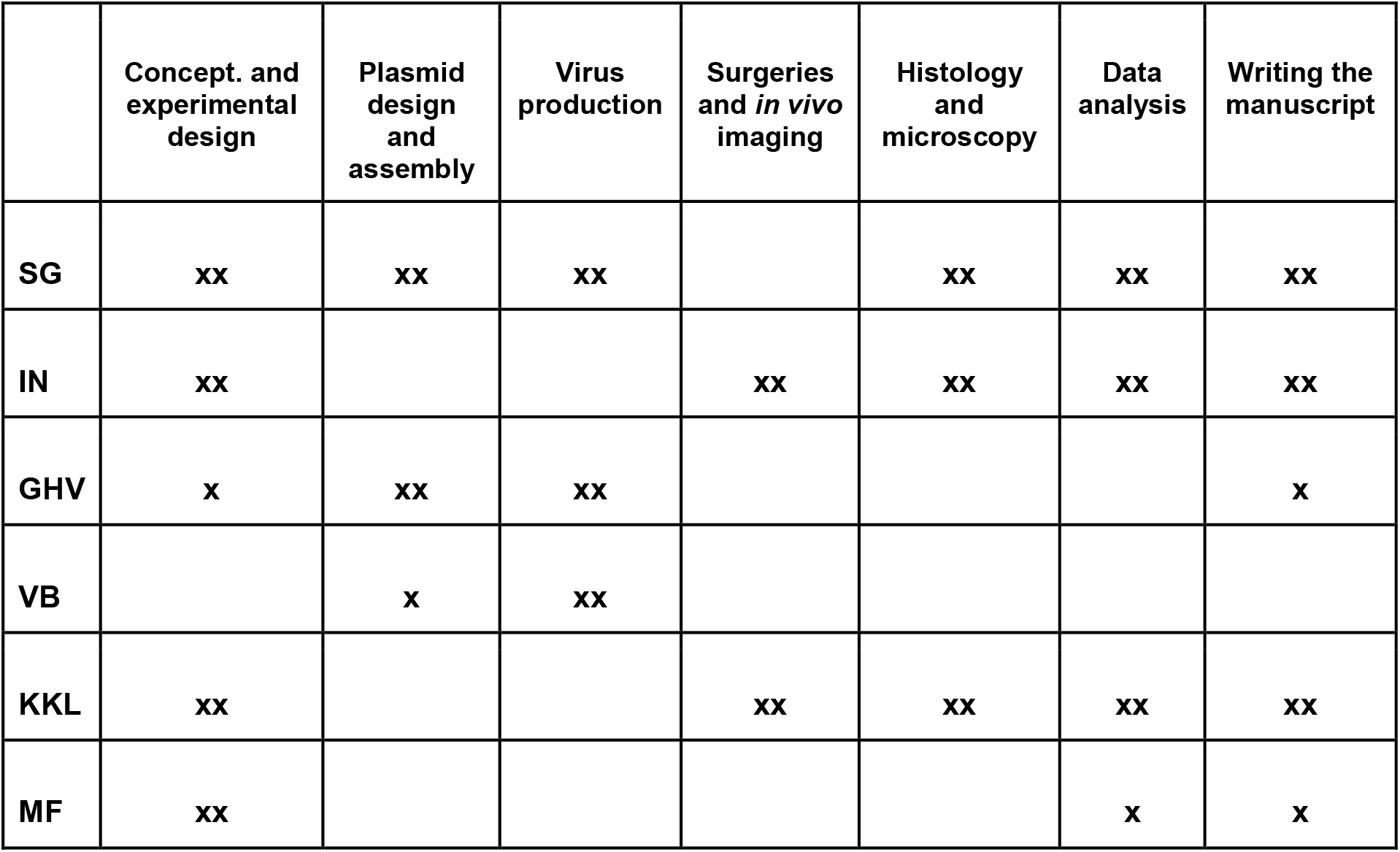

All authors have read the final version of the manuscript. The authors report no conflicts of interest.

## Acknowledgements

We thank Dr. Ane Charlotte Chistensen and Dr. Jennifer L. Hazen for assistance with establishment of the RO injection technique, Paul Johannes Helm for assistance with assembly of the two-photon microscope and the Instrumentation Lab at the Dept of Bioscience. Imaging of histological sections was performed at the NorMIC microscopy platform at the Dept. of Bioscience, University of Oslo.

## Supplementary Information

The supplementary information contains supplementary table S1 and supplementary figures S1-3.

In addition, 7 supplementary videos are attached to the manuscript.

**Figure S1:**
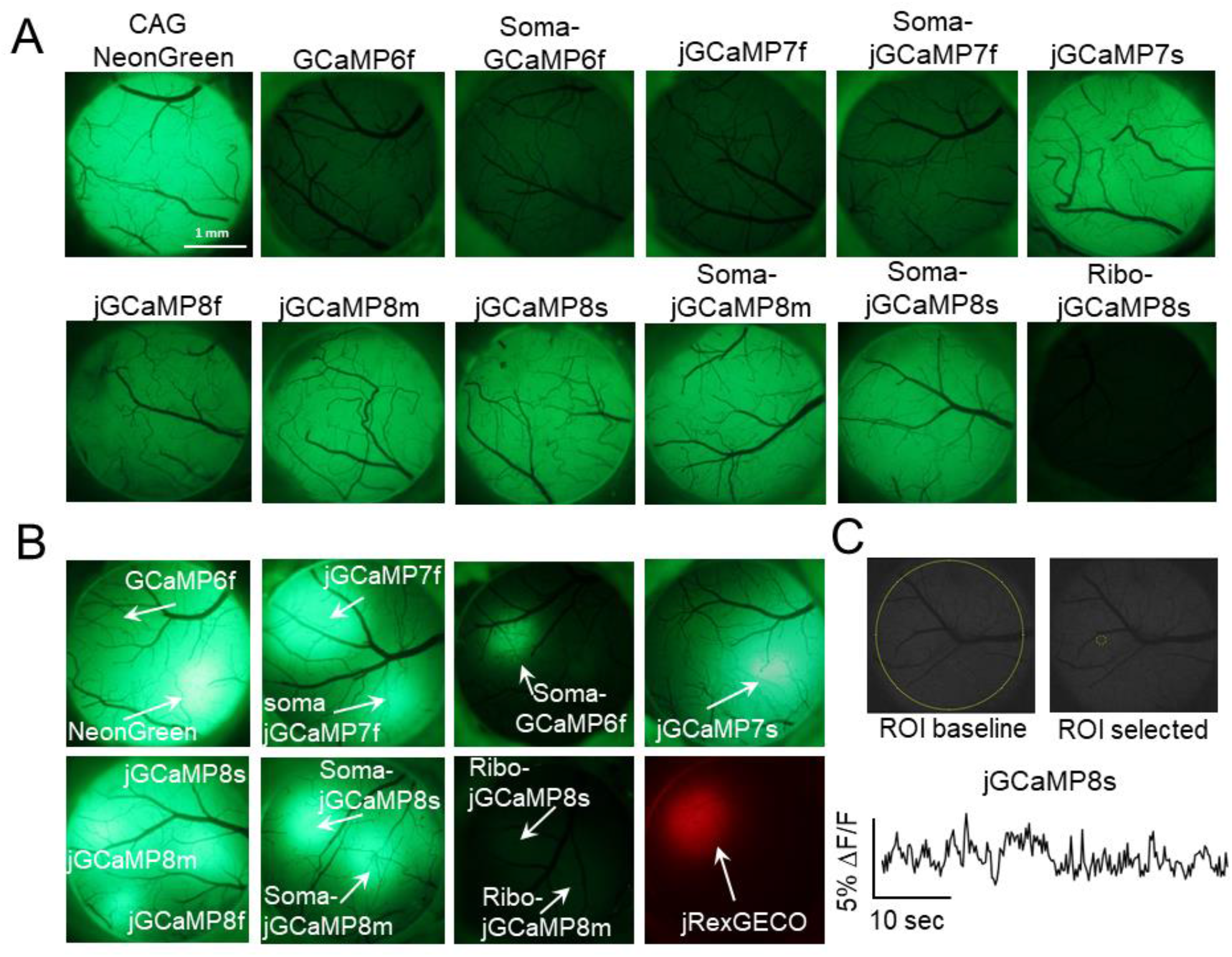
Overview of GECI screening. Cranial windows were made over visual cortex and monitored by widefield fluorescence microscopy. **A** shows RO injected animals, while **B** shows local virus injections. **C**: Widefield imaging of spontaneous cortical activity did not indicate seizures from brainwide GECI expression. Example of a region of interest and Ca2+ trace.

**Figure S2:**
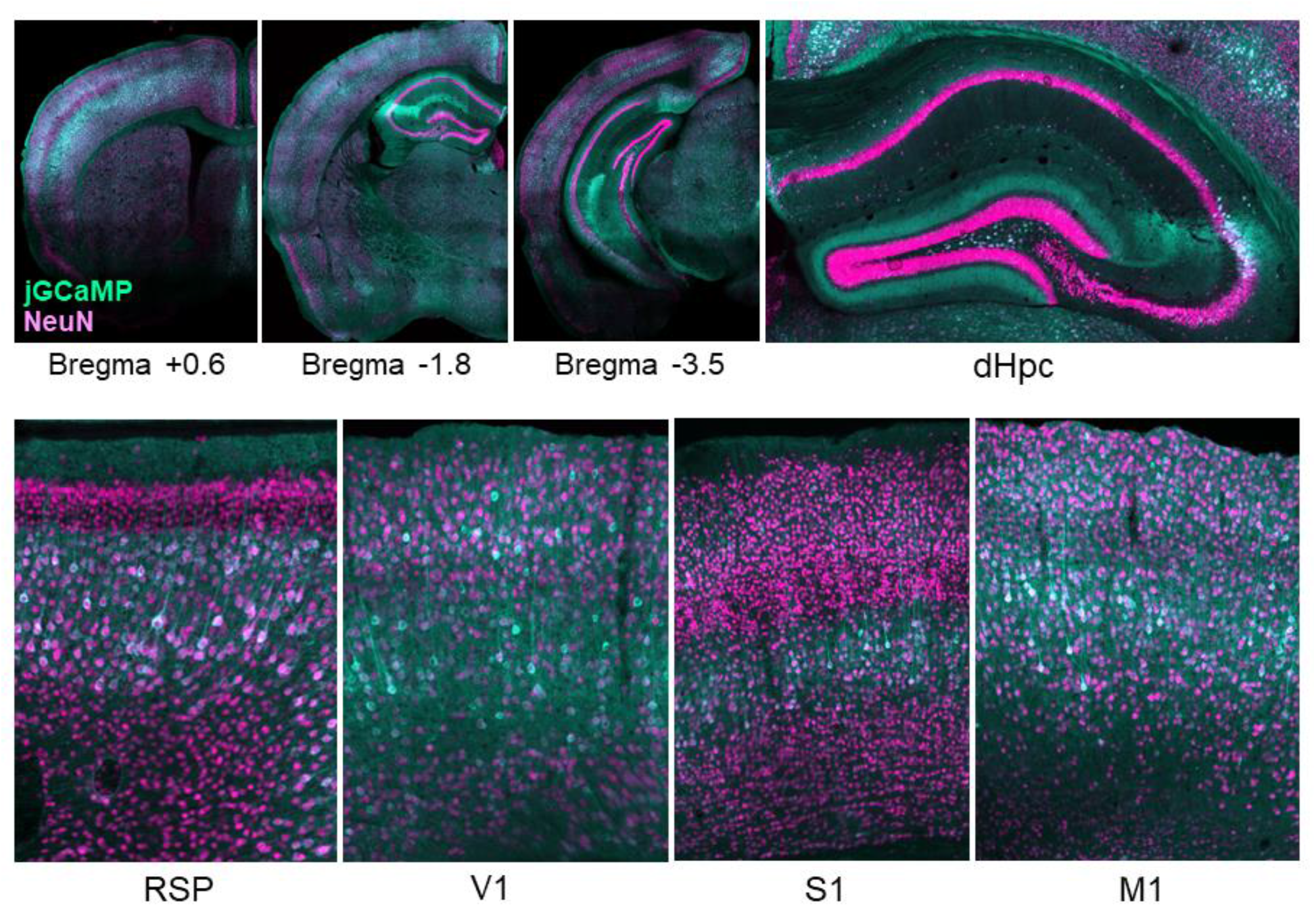
Examples of post-mortem histological samples from different brain areas, stained with GFP and the neuronal marker NeuN.

**Figure S3:**
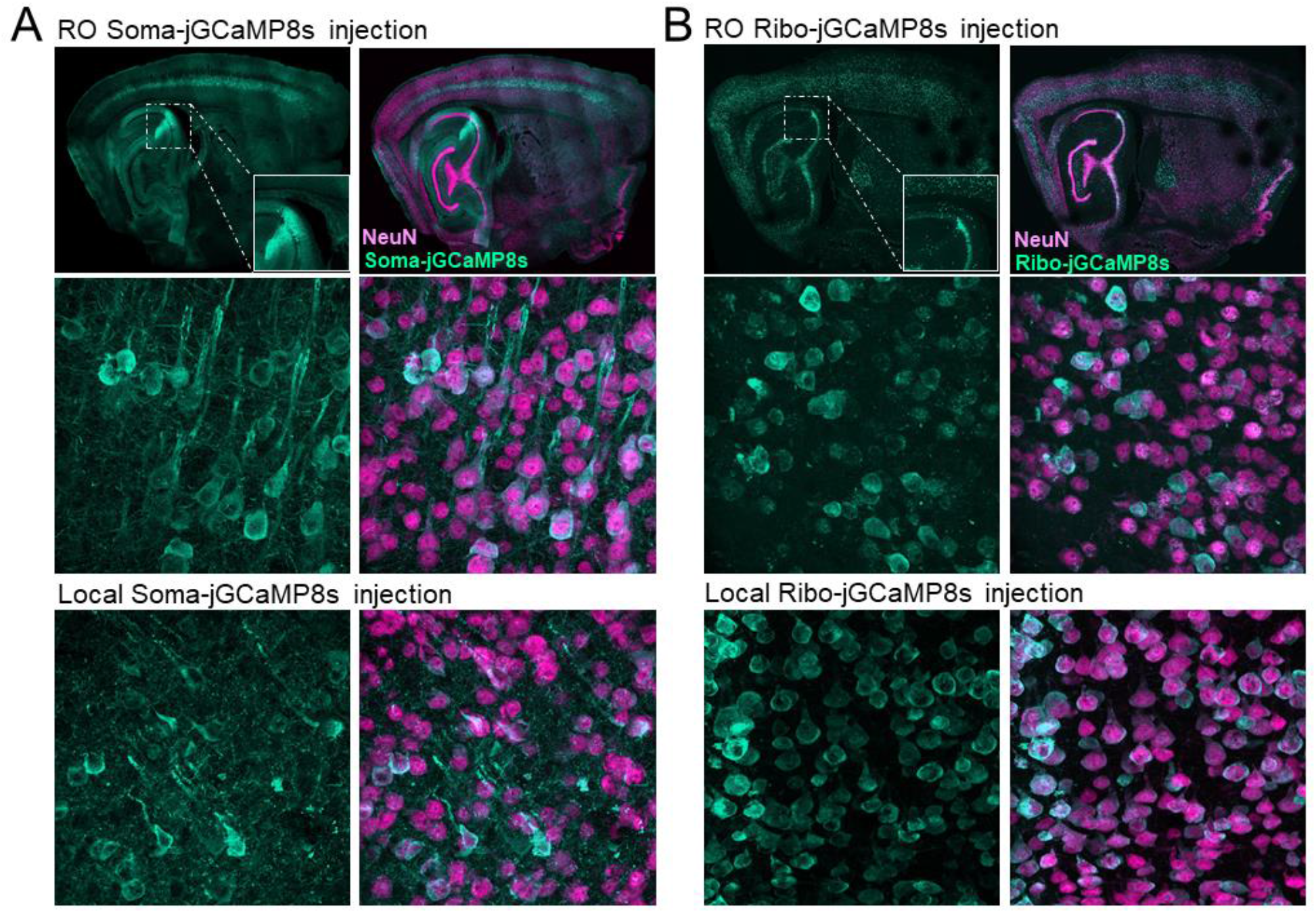
Histological verification of expression from soma-targeted expression of jGCaMP8. **A**: EE-RR soma targeting using RO or local virus injections. Upper panel shows a saggital section of a mouse brain with hippocampal region CA2 highlighted. Lower panels show high-resolution images from primary visual cortex from both systemic and local virus injections. **B**: Same as for A, but using ribosome-tethered jGCaMP8s.

**Table S1:**
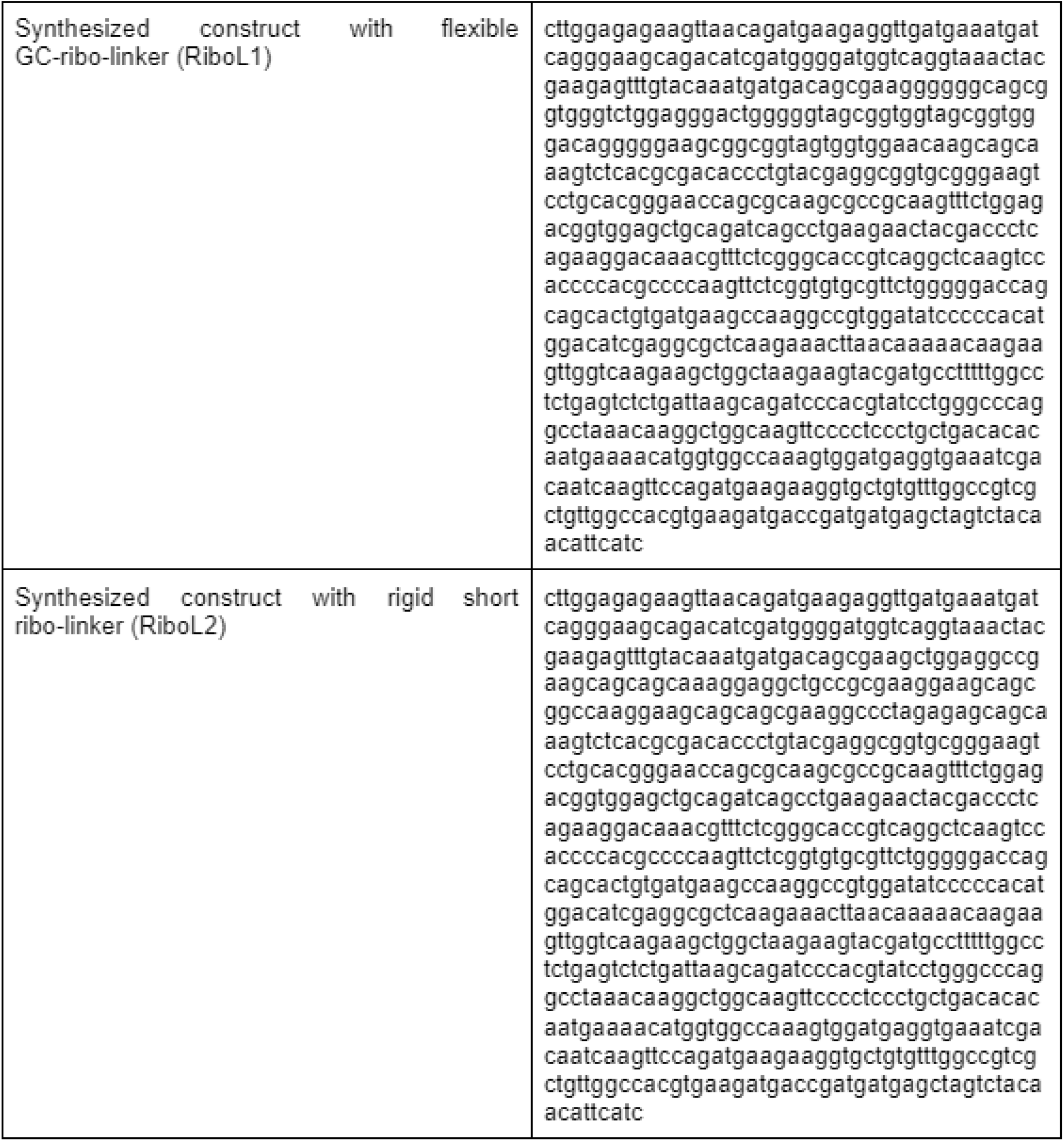
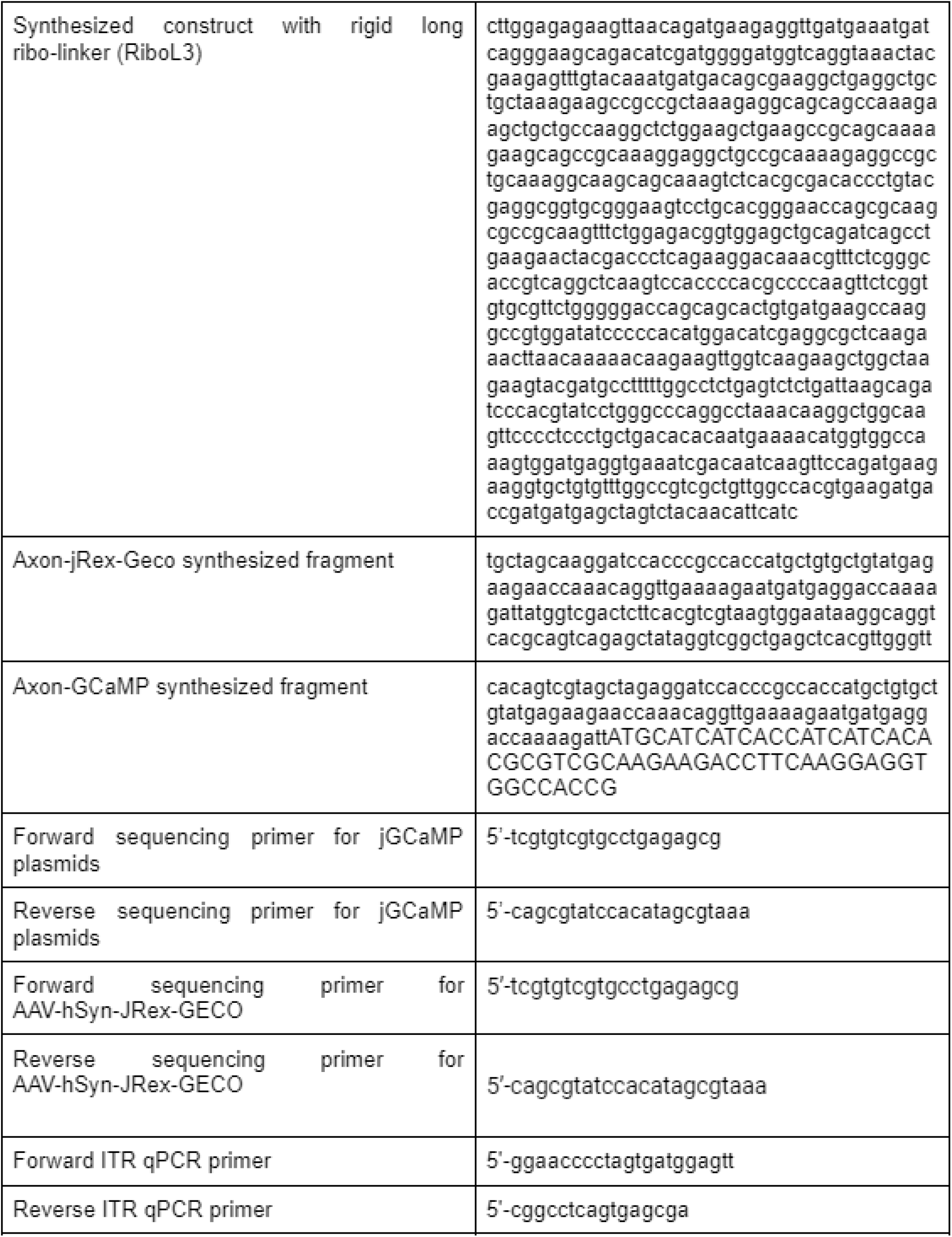
Primers and DNA sequences, 5’ to 3’:

